# stDyer-image improves clustering analysis of spatially resolved transcriptomics and proteomics with morphological images

**DOI:** 10.1101/2025.02.07.634992

**Authors:** Ke Xu, Xin Maizie Zhou, Lu Zhang

## Abstract

Spatially resolved transcriptomics (SRT) and spatially resolved proteomics (SRP) data enable the study of gene expression and protein abundances within their precise spatial and cellular contexts in tissues. Certain SRT and SRP tech-nologies also capture corresponding morphology images, adding another layer of valuable information. However, few existing methods developed for SRT data effectively leverage these supplementary images to enhance clustering performance. Here, we introduce stDyer-image, an end-to-end deep learning framework designed for clustering for SRT and SRP datasets with images. Unlike existing methods that utilize images to complement gene expression data, stDyer-image directly links image features to cluster labels. This approach draws inspiration from pathologists, who can visually identify specific cell types or tumor regions from morphological images without relying on gene expression or protein abundances. Benchmarks against state-of-the-art tools demonstrate that stDyer-image achieves superior performance in clustering. Moreover, it is capable of handling large-scale datasets across diverse technologies, making it a versatile and powerful tool for spatial omics analysis.

## Introduction

Spatially resolved transcriptomic (SRT) and spatially resolved proteomics (SRP) measure spatial coordinates in addition to gene expression data or proteomics data, enabling the investigation of biological dynamics within their spatial contexts. More-over, some SRT and SRP technologies [1–5] can simultaneously capture spatially annotated images, further enhancing the resolution and interpretability of the data. A critical task in SRT data analysis is identifying spatial domains where units exhibit similar gene expression patterns by considering the relationship between a target unit (either spots or cells, depending on the SRT technologies) and its spatial neighbors. Similarly, identifying cell types in SRT and SRP data is essential to facilitate down-stream biological analysis. To address these tasks, various clustering methods [6–12] have been developed to incorporate spatial coordinates into the clustering process. However, only a few of them [8–12] utilize additional image information to further enhance clustering performance and improve biological insights.

SpaGCN [8] transforms morphological features of the units into artificial coordinates and integrates them with spatial coordinates to construct a graph that serves as input for its clustering algorithm. stLearn [9] employs matrix multiplication to combine a gene expression correlation matrix and a morphological similarity matrix for imputation. The imputed matrix is subsequently used to perform clustering analysis. SiGra [10] extracts image patches, flattens them into vectors, and reconstructs gene expression data from input to obtain augmented embeddings for clustering. DeepST [11] incorporates a morphological similarity matrix, a gene expression correlation matrix, and a spatial adjacency matrix to enhance gene expression data and generate embeddings for clustering. MUSE [12] employs a reconstruction loss to facilitate the mutual reconstruction of image and transcriptomic modalities, while a self-supervision loss ensures consistency of relative distance relationships between units across individual and joint modalities, optimizing embeddings for clustering analysis.

Although these methods have managed to combine transcriptome and image modalities for clustering analysis, several limitations still remain. First, these methods focus on either reconstructing one modality from the other or improving graph construction to derive informative embeddings with an independent clustering step. This decoupled process may lead to suboptimal performance as clustering is not performed in an end-to-end manner. Second, these methods often support only one or two specific technologies, and applying them to custom datasets typically requires modifications or a thorough understanding of their source code due to insufficient documentation on input format specifications for the image modality. Third, existing tools that utilize the image modality with GPU acceleration are not scalable to large-scale datasets due to graphics memory limitation.

To address these limitations, we introduce stDyer-image, an end-to-end, scalable clustering approach for spatially resolved omics data. Unlike existing methods, stDyer-image directly associates the image modality with predicted labels to enhance clustering performance. This design draws inspiration from the practice of pathologists or trained doctors who can visually identify specific cell types or tumor regions by examining images. Furthermore, stDyer-image employs a Gaussian Mixture Variational AutoEncoder (GMVAE), which directly optimizes predicted labels using objective functions that incorporate image similarities, further improving clustering accuracy and robustness. Additionally, stDyer-image is compatible with a wide range of spatially resolved omics technologies, including CosMx, Stereo-seq, 10x Xenium, 10x Visium HD, and CODEX. Moreover, stDyer-image incorporates a mini-batch neighbor sampling strategy and supports multi-GPU training, enabling its application to large-scale datasets. We compared stDyer-image with eight state-of-the-art tools across five different technologies, demonstrating its superior performance and wide applicability.

## Results

### Workflow of stDyer-image

stDyer-image builds on the GMVAE [13], integrating it with Graph ATtention network (GAT) [14] to generate embeddings and cluster labels, similar to stDyer [15]. The GMVAE processes inputs such as gene expression profiles or protein abundance profiles, along with a KNN graph (Figure 1). The KNN graph can be constructed based on spatial coordinates, gene expression profiles, or protein abundance matrix, depending on whether the task involves identifying cell types or delineating spatial domains. The major difference between stDyer-image and stDyer is that stDyer-image incorporates image patches into an image-related loss function to improve clustering performance.

**Fig. 1:**
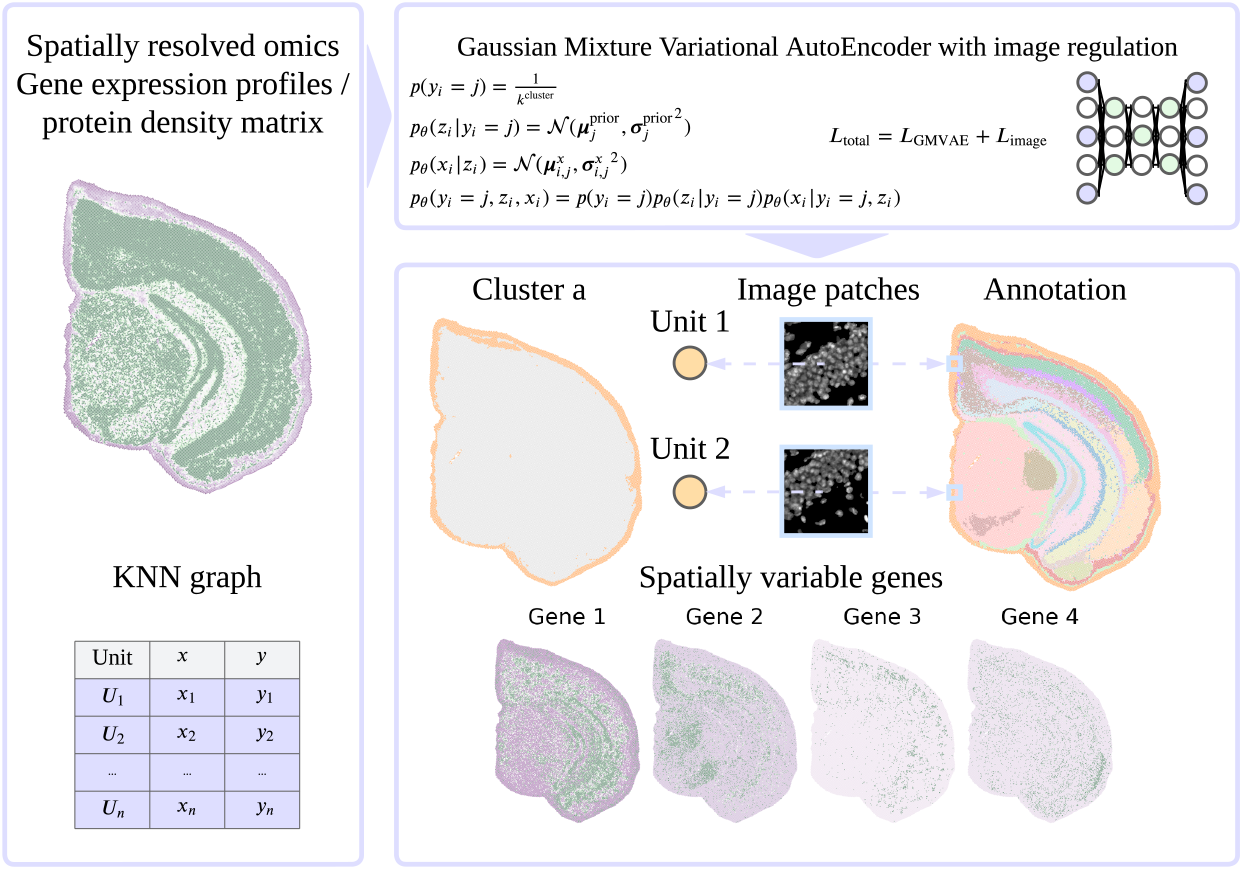
The workflow of stDyer-image. stDyer-image takes as input the gene expression profiles or protein abundance matrix along with a KNN graph. The GMVAE, incorporating a GAT, generates embeddings and cluster labels, which are optimized using loss functions that account for gene expression, protein abundance, and image data. The loss function involving gene expression profiles or protein abundance profiles maximizes the log-likelihood of the marginal distribution for modeling these profiles. This encourages neighboring units to have similar embeddings and cluster labels, relying solely on gene expression profiles or protein abundance profiles combined with spatial coordinates. The image-related loss function encourages units with similar image embeddings to share similar soft cluster labels. Additionally, stDyer-image can identify spatially variable genes using integrated gradient analysis.

The loss functions of stDyer-image comprise the original function from stDyer and an additional image-related loss function. The image-related loss function involves the soft probabilities of predicted cluster labels and the cosine similarity between the soft cluster labels of a unit and its image neighbors. Image neighbors are defined as units with similar image patches to the target unit. This image-related loss establishes a connection between the soft cluster label of a unit and its image neighbors, encouraging units with similar image patches to share similar soft cluster labels. The loss function from stDyer includes the log-likelihood of the marginal distribution for modeling gene expression data or protein abundance data, as well as the reconstruction loss of these data. This component promotes similar embeddings and soft cluster labels among neighboring units, relying solely on gene expression or protein abundance data. By integrating the image-related loss, stDyer-image further enhances clustering performance by incorporating image-derived information.

### stDyer-image identifies tumor on the NSCLC dataset from CosMx technology

We evaluated the performance of stDyer-image on a non-small cell lung cancer (NSCLC) dataset [1] generated using CosMx technology. This dataset consists of 20 slices (Figure 2a) and their associated images (Figure 2b), resulting in a total of 87,606 units. These slices are annotated with eight cell types (Figure 2a) and arranged in a 5×4 grid to accommodate a large area of tissue. We benchmarked stDyer-image against eight state-of-the-art methods for clustering analysis on spatial transcriptomics data: BayesSpace, CellCharter, stDyer, SpaGCN, stLearn, SiGra, DeepST, and MUSE, using adjusted rand index (ARI; **Methods**) as the evaluation metric. Among these, BayesSpace, CellCharter, and stDyer are scalable to large datasets but are incapable of using images, whereas the remaining methods are capable of utilizing images for clustering analysis. Importantly, stDyer-image, stDyer, BayesSpace, CellCharter, and SiGra can analyze all 20 slices simultaneously and provide consistent predictions across slices, while the other methods are limited to processing one slice at a time. We reported the single-slice ARI scores of each method in the box plot (Figure 2c) and observed that stDyer-image performed the best on the NSCLC dataset with an average ARI of 0.561 across all 20 slices (Figure 2d). stDyer-image demonstrated superior performance in predicting tumor regions while stDyer, BayesSpace, and CellCharter tended to split them into multiple parts. While SiGra identified the entire tumor regions, it failed to accurately identify neutrophil cells located in proximity. SpaGCN, stLearn, and DeepST struggled to produce consistent cluster labels across slices.

**Fig. 2:**
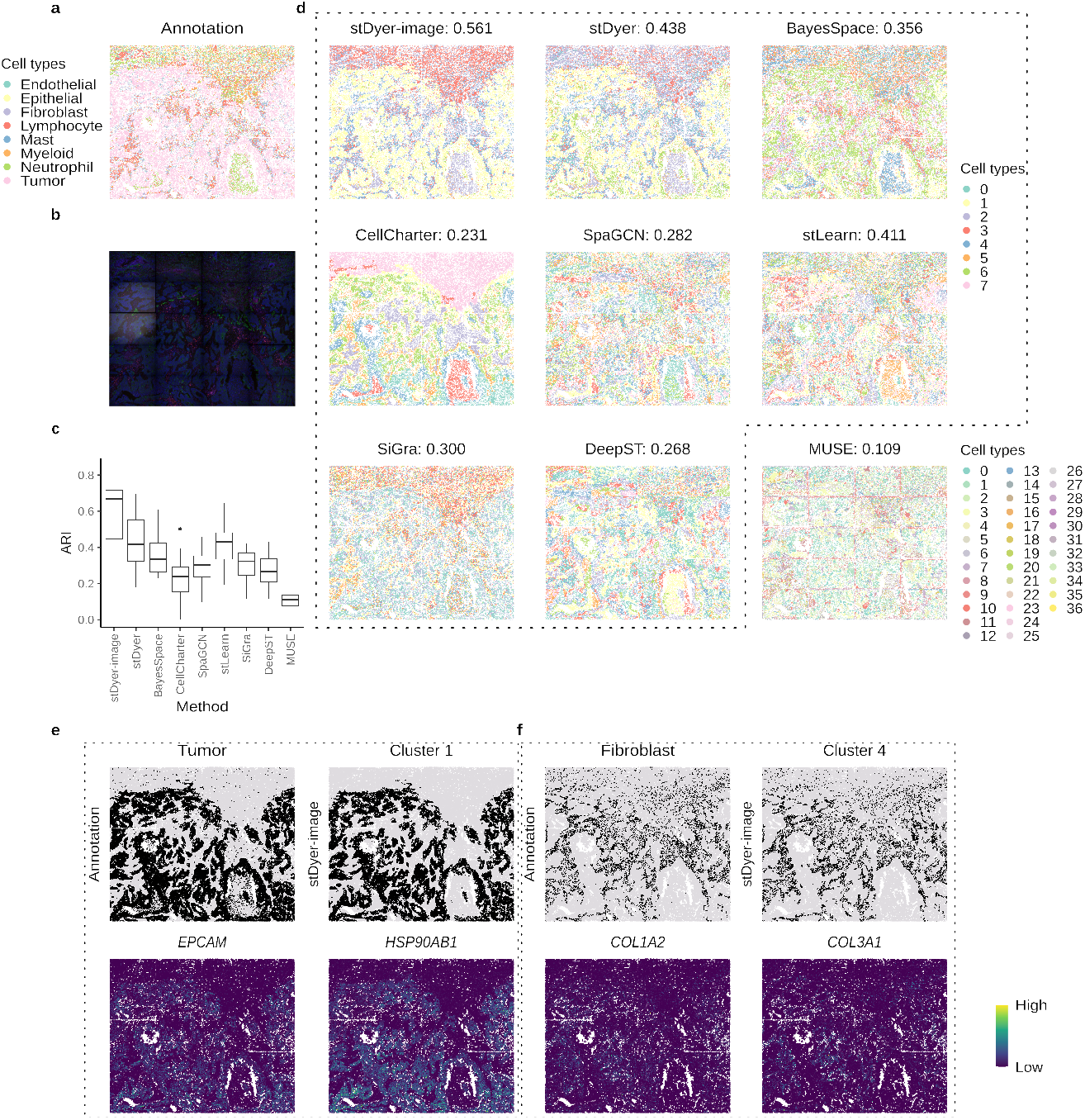
Performance of stDyer-image on a human non-small cell lung cancer dataset from CosMx technology. (**a**) Visualization of the annotation for the human NSCLC dataset. (**b**) Morphology image of the human NSCLC dataset. (**c**) Boxplot of ARI scores for nine methods evaluated across 20 slices of the human NSCLC dataset. (**d**) Visualization and the average ARI scores across 20 slices for cell type clustering using different methods on the human NSCLC dataset. (**e**) Visualization of the annotation for tumor regions, the prediction of cluster 1 by stDyer-image, and two selected SVGs for cluster 1. (**f**) Visualization of the annotation for fibroblast regions, the prediction of cluster 4 by stDyer-image, and two selected SVGs for cluster 4.

MUSE did not accept a fixed cluster number as input and produced an excessive number of clusters. Furthermore, MUSE performed poorly on slice boundaries.

To explore whether the domains identified by stDyer-image facilitate the detection of biologically relevant biomarkers, we ranked the top 50 spatially variable genes (SVGs) with the highest integrated gradient (IG) values for each cluster (**Methods**). For visualization purposes, we selected two genes that strongly aligned with cluster 1 and 4, respectively (Figure 2e and f). For instance, cluster 1, which corresponded to the tumor region in the annotation (Figure 2e), included *EPCAM* as one of its SVGs. The protein EPCAM is known to be widely present in NSCLC tumor regions but not in normal Lung tissue [16]. Additionally, another SVG for cluster1, *HSP90AB1*, was highly expressed in NSCLC tumor tissue and is associated with poor prognosis in lung adenocarcinoma patients [17]. Cluster 4, corresponding to fibroblasts in the annotation (Figure 2f), included *COL1A2* and *COL3A1* as highly expressed SVGs. The cluster associated with these genes was classified as matrix cancer-associated fibroblast by [18].

### stDyer-image recognizes smooth laminar layers on the mouse brain dataset from Stereo-seq technology

The mouse brain dataset [2], generating using Stereo-seq technology, comprises 38,746 units and 23,905 genes, with 20 annotated spatial domains (Figure 3a and b). We benchmarked stDyer-image against stDyer, BayesSpace, CellCharter, SpaGCN and stLearn. Among these methods, stDyer-image achieved the highest ARI score of 0.517 (Figure 3c). Other methods (SiGra, DeepST, and MUSE) failed to run on this large dataset due to out of memory (OOM) errors. stDyer-image produced more spatially coherent layer predictions compared to stDyer, BayesSpace, SpaGCN, and stLearn. While CellCharter achieved even smoother predictions, it failed to identify thin layers such as the Dentate gyrus. We performed a similar SVG analysis, identifying SVGs associated with specific clusters based on their IG values (Figure 3d and e). For example, the Dentate gyrus was identified as cluster 1 (Figure 3d), with the SVG *Prox1* (Figure 3d) playing a crucial role in the maintenance and maturation of Dentate gyrus granule cells [19, 20]. Additionally, the SVG *C1ql2* was found to be highly expressed in the Dentate gyrus [21]. In the Ventral tegmental area, the SVG *Slc6a3* (Figure 3e), encoding the dopamine transporter, was associated with the rewarding function of this region [22]. Another SVG, *Th* (Figure 3e), a known marker of dopaminergic neurons [22], was also identified to be highly expressed in the Ventral tegmental area.

**Fig. 3:**
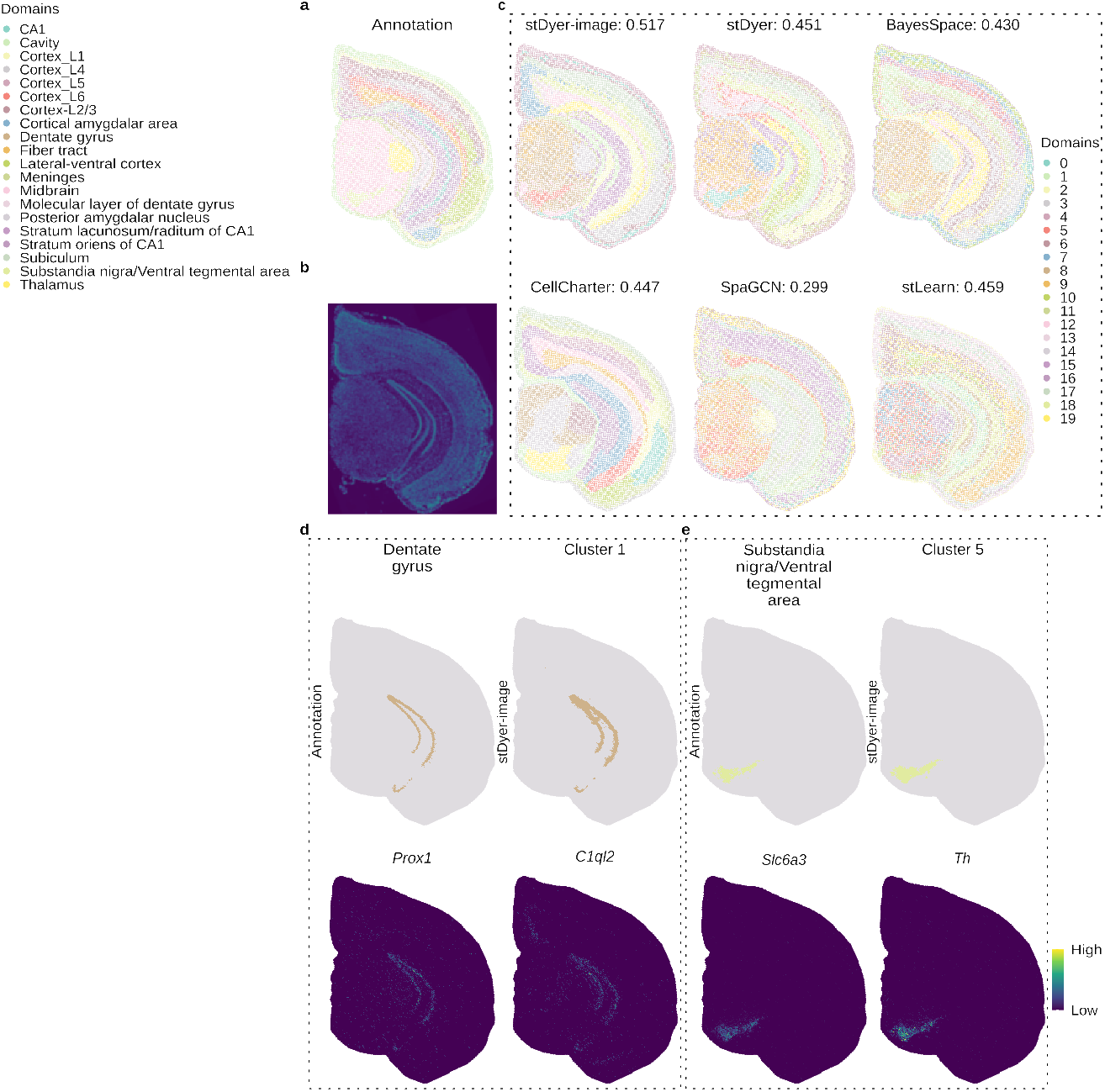
Performance of stDyer-image on a mouse brain dataset from Stereo-seq technology. (**a**) Visualization of the annotation for mouse brain tissue. (**b**) Morphology image of mouse brain tissue. (**c**) Visualization and ARI scores of various methods for spatial domain clustering on mouse brain tissue. (**d**) Visualization of the annotation for the Dentate gyrus, the prediction of cluster 1 by stDyer-image, and the two selected SVGs for cluster 1. (**e**) Visualization of the annotation for Substandia nigra/Ventral tegmental area, the prediction of cluster 5 by stDyer-image, and the two selected SVGs for cluster 5.

### stDyer-image detects proliferative invasive tumor on a human breast cancer dataset from 10x Xenium technology

We also analyzed a human breast cancer dataset [3] from 10x Xenium technology, consisting of 159,226 units and 313 genes, annotated with 19 cell types (Figure 4a and b). We benchmarked stDyer-image against stDyer, BayesSpace, CellCharter, SpaGCN and stLearn. stDyer-image achieved the highest ARI score of 0.516 (Figure 4c). SiGra, DeepST, and MUSE still failed to process this large dataset due to OOM errors. Further comparisons were made across the Stromal and Invasive Tumor clusters, which represent the two largest annotated clusters in the dataset (41,422 units and 34,374 units, respectively, out of 159,226 units). stDyer-image, stDyer, BayesSpace, and SpaGCN identified the most extensive region for Stromal. While stLearn performed well overall, it introduced segmentation artifacts At the right center. CellCharter, on the other hand, identified more units belonging to Stromal in the central region but divided the cluster into smaller clusters at the top and bottom. For Invasive Tumor, only stDyer-image and stLearn predicted 29,347 and 26,304 units in the cluster 2 and 13, respectively. Other methods predicted units fewer than 15,000 units in their largest cluster matched with Invasive Tumor.

**Fig. 4:**
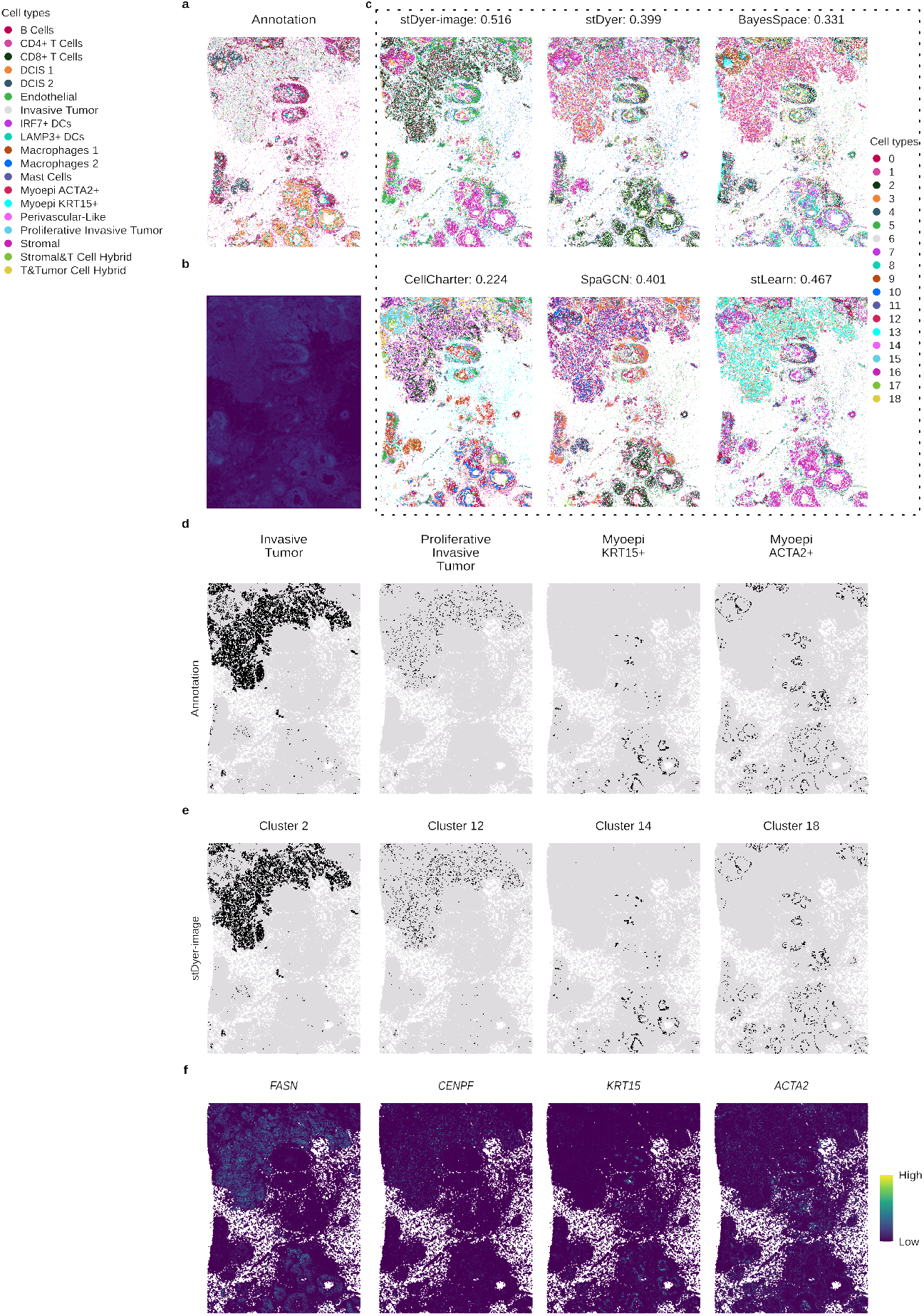
Performance of stDyer-image on a human breast cancer dataset from 10x Xenium technology. (**a**) Visualization of the annotation for the human breast cancer dataset. (**b**) Morphology image of the human breast cancer dataset. (**c**) Visualization and ARI scores of different methods for cell type clustering on the human breast cancer dataset. (**d**) Visualization of the annotation for Invasive Tumor, Proliferative Invasive Tumor, Myoepi KRT15+, and Myoepi ACTA2+. (**e**) Visualization of predictions for 8 clusters 2, 12, 14, and 18 from stDyer-image. (**f**) Visualization of SVG associated with clusters 2, 12, 14, and 18 from stDyer-image.

We performed IG analysis and found that the Invasive Tumor (Figure 4d) corresponded to cluster 2 (Figure 4e), which was associated with the SVG *FASN* (Figure 4f). *FASN* is known to be overexpressed in human breast cancer, serving as a marker of poor prognosis [23]. It mediates changes in certain fatty acids that promote tumor migration [24]. Similarly, *CENPF*, the SVG for cluster 12, was found to be associated with Proliferative Invasive Tumor. Overexpression of this gene has been linked to tumor bone metastasis in breast cancer [25] and can result in chromosomal instability [26], and it is also indicative of poor prognosis [25, 26]. Additionally, the SVGs *KRT15* and *ACTA2* were identified as being associated with clusters 14 and 18, corresponding to Myoepi KRT15+ and Myoepi ACTA2+ regions, respectively, as indicated in the annotations.

### stDyer-image identifies glands structure on the human colon cancer dataset from 10x Visium HD technology

We downloaded a human colon cancer dataset [4] (Figure 5a and b) from a tutorial on 10x Visium HD technology to evaluate the performance of benchmarked methods. This technology utilizes 2*µm ×* 2*µm* squares to measure tissue, comprising 8,731,400 units and 18,085 genes. Due to the dataset’s large size, direct analysis of the original units was impractical. To address this, the dataset was aggregated into larger binned units of either 8*µm ×* 8*µm* squares or 16*µm ×* 16*µm* squares, resulting in 545,913 and 137,051 binned units, respectively. For our evaluation, we used the 16*µm ×* 16*µm* binned units since fewer methods were capable of handling the larger dataset. Among these benchmarked methods, only stDyer-image, stDyer, SpaGCN, stLearn, and CellCharter successfully analyzed this dataset. BayesSpace failed due to a limitation in its dependent package, which could not accept a too large matrix. Similarly, SiGra, DeepST, and MUSE could not analyze this dataset due to OOM errors. This dataset did not have annotations for all units, so we first performed K-Means clustering and computed corresponding Silhouette scores for different numbers of clusters. The Silhouette scores for 10, 15, 20, 25, and 30 clusters were 0.248, 0.265, 0.213, 0.189, and 0.191, respectively. Based on these results, we selected 15 clusters for benchmarking all methods. Overall, all methods successfully identified the general spatial structure (Figure 5b). Compared to all other methods, stDyer-image uniquely distinguished the region in the middle bottom part as the 2nd, 6th, and 12th clusters (Figure 5b), which aligned with the morphological image (Figure 5a).

**Fig. 5:**
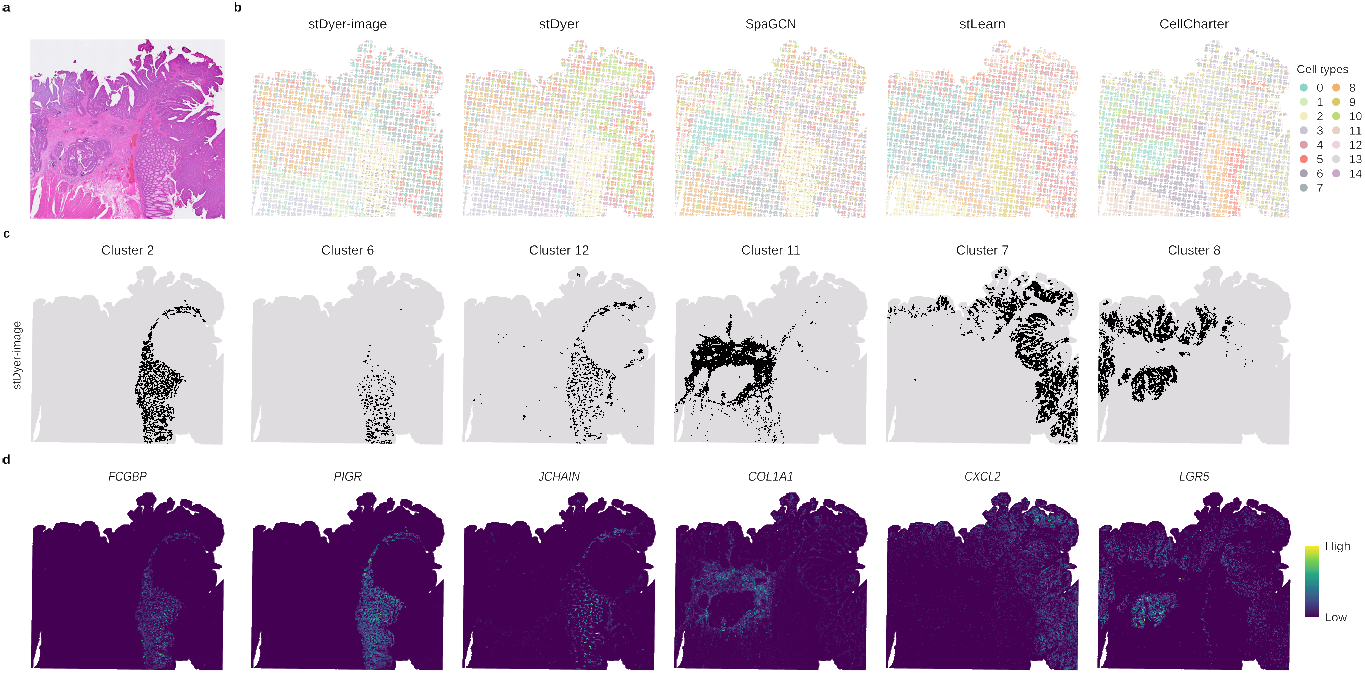
Performance of stDyer-image on a human colon cancer dataset from 10x Visium HD technology. (**a**) Morphology image of the human colon cancer dataset. (**b**) Visualization and ARI scores of different methods for cell type clustering on the human colon cancer dataset. (**c**) Visualization of predictions for clusters 2, 6, 12, 11, 7 and 8 from stDyer-image. (**d**) Visualization of SVGs associated with clusters 2, 6, 12, 11, 7, and 8 from stDyer-image.

We performed IG analysis and identified the SVG *FCGBP*, associated with cluster 2 (Figure 5c), as being highly expressed (Figure 5d). *FCGBP* is a known marker of goblet cells within this cluster and encodes protein FCGBP, which is secreted into the mucus [27]. stDyer-image further distinguished cluster 6, which occupied the inner layer compared to the outer layer composed of goblet cells. The SVG *PIGR*, associated with this cluster, is a known marker of goblet cells and enterocytes [4]. Furthermore, the external region surrounding goblet cells was identified as cluster 12, with the SVG *JCHAIN* serving as its marker. *JCHAIN* is a recognized marker of plasma cells [28]. stDyer-image also identified fibroblast regions and associated SVG *COL1A1*, which is known to induce fibroblasts to transform into tumor-associated fibroblasts [29]. Additionally, tumor regions were identified by stDyer-image, indicated by the SVGs *CXCL2* and *LGR5. CXCL2* can promote the tumor cell proliferation [30], while *LGR5* serves as surface markers of cancer stem cells in colorectal cancer [31].

### stDyer-image deciphers clear laminar structures on the human intestine dataset from CODEX technology

Given the similarity in format between spatially resolved proteomics data and spatial transcriptomics data, we applied stDyer-image to a human intestine dataset [5] from CODEX technology. This dataset contained 64 slices from 8 donors, with each slice characterized by measurements of 47 proteins. For each donor, 8 slices were included, with 4 from the colon and 4 from the small intestine. These slices are non-adjacent and spatially separated. We first benchmarked stDyer-image on two slices, one from the colon (Figure 6a and b) and one from the small intestine (Figure 6d and e). For the colon tissue, stDyer-image achieved an ARI score of 0.400 (Figure 6c), outperforming the second-best method, stLearn (ARI=0.293). Compared to stLearn, stDyer-image identified more heterogeneity in the leftmost layers. For example, cluster 1 predicted by stDyer-image corresponded to a region that aligned with Adaptive Immune Enriched domain. Additionally, stDyer-image uniquely identified the distinct left and right regions within the Smooth Muscle domain, whereas other methods erroneously mixed the left region with the Stroma domain (Figure 6f). For the small intestine tissue, stDyer-image achieved an ARI score of 0.361, outperforming the second-best method stDyer (ARI=0.036). All other methods failed to delineate continuous domains and instead mixed different domains together. DeepST and MUSE failed to process this slice due to OOM errors. We further benchmarked stDyer-image on all 8 slices from donor B008. SpaGCN was incompatible with datasets because it requires at least 50 features to perform PCA internally, while CellCharter could not be applied because the dataset provided normalized values rather than raw counts. SiGra, DeepST, and MUSE failed due to OOM errors. stDyer-image achieved the highest average ARI score of 0.346, outperforming stDyer (ARI=0.157), stLearn (ARI=0.152), and BayesSpace (ARI=0.082) (Figure 6g).

**Fig. 6:**
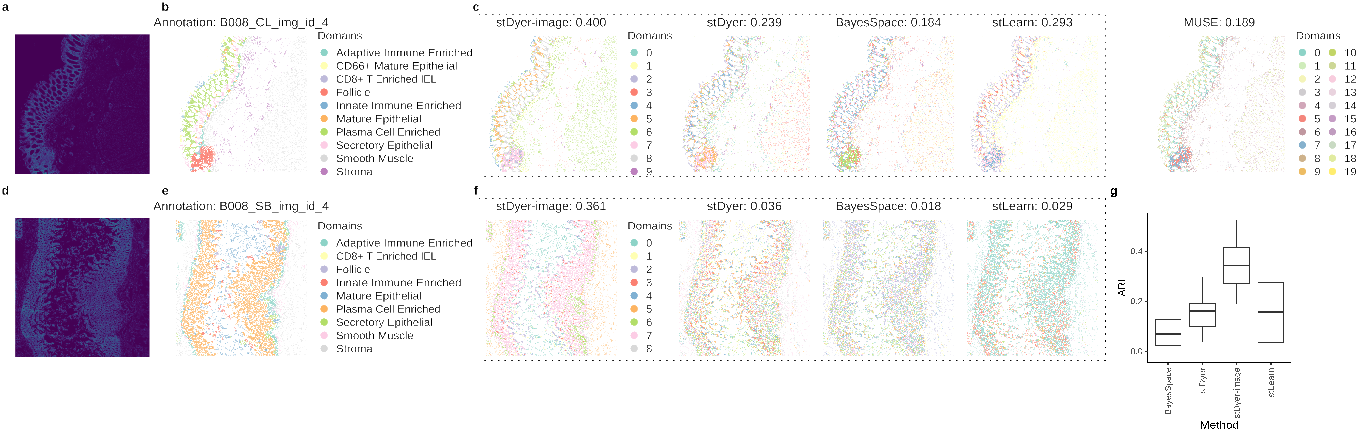
Performance of stDyer-image on a human intestine dataset from CODEX technology. (**a**) Morphology image of the 4th slice from colon tissue of the B008 donor. **(b)**Visualization of the annotation for the 4th slice from colon tissue of the B008 donor. (**C)**Visualization and ARI scores of different methods for spatial domain clustering on the 4th slice from colon tissue of B008 donor. (**d**) Morphology image of the 4th slice from small bowel tissue of the B008 donor. (**e**) Visualization of the annotation for the 4th slice from small bowel tissue of the B008 donor. (**f**) Visualization and ARI scores of different methods for spatial domain clustering on the 4th slice from small bowel tissue of the B008 donor. (**g**) Boxplot of ARI scores for four methods evaluated across all eight slices from the B008 donor.

## Discussion

During benchmarking of existing clustering methods designed for spatially resolved omics data, we observed that the scalability of most methods is limited. Some are constrained by the available GPU memory since they lack a mini-batch strategy to handle large-scale datasets. Others suffer from inefficient optimization, with excessive IO operations related to image processing that significantly slow down clustering analysis. In contrast, stDyer-image is designed to be more memory-efficient, using a mini-batch strategy to process large-scale datasets. Additionally, IO operations are minimized in stDyer-image to speed up clustering analysis when images are used as input, especially when temporary file storage relies on slow-speed devices such as hard disk drives.

stDyer-image is compatible with data from various technologies, unlike other methods that utilize images but are restricted to one or two technologies. Running those methods on unsupported datasets often requires modification to their code or the dataset structure. This process can be challenging due to a lack of detailed documentation on the required input format. In contrast, we have provided comprehensive descriptions of the input format, facilitating the use of stDyer-image on customized datasets. Additionally, a user-friendly interface allows users to run stDyer-image on their datasets with just a few lines of code. Unlike many other methods that require raw counts as input, stDyer-image can also process normalized data, offering greater flexibility and ease of use. Furthermore, stDyer-image supports multi-GPU acceleration, significantly enhancing its capability to analyze large-scale datasets efficiently. These features make stDyer-image a robust and accessible tool for spatial omics data analysis.

## Methods

### Data preprocessing

An SRT or SRP dataset with images should include three major components: (1) a gene expression matrix or protein abundance matrix with *m* genes or proteins for *n* units (***X*** *∈* ℝ^*n×m*^); (2) spatial coordinates indicating the locations of units; and (3) an image capturing the texture and density of units. Preprocessing of the gene expression matrix in SRT datasets was performed following standard conventions. Specifically, unless stated otherwise, we used Scanpy [32] as follows: 1. removed feature names starting with “NegPrb” for the non-small-cell lung cancer dataset from CosMx; 2. removed unlabeled units in the human breast cancer dataset from Xenium and units with spatial coordinates outside the image for all datasets; 3. filtered units with zero RNA counts using [scanpy.pp.filter cells(min counts=1)] and genes with zero RNA counts using [scanpy.pp.filter genes(min counts=1)]; 4. selected the top 3,000 highly variable genes using [scanpy.pp.highly variable genes]; 5. performed library size normalization using [scanpy.pp.normalize total]; 6. applied logarithmic transformation with [scanpy.pp.log1p]; 7. standardized gene expression values to z-score using [scanpy.pp.scale].

For images, we selected the highest-resolution image (typically with the .tif or.tiff suffix) for each dataset and cropped them into 250×250-pixel patches. We determined that a patch size of 250×250 pixels adequately represents the local environment of a cell and its neighboring cells (Figure 1). For single-channel images, we converted them into three channels using [numpy.broadcast to] [33] to facilitate feature extraction. For the mouse brain dataset from Stereo-seq, we employed a modified version of the “read bgi agg” function from the spateo package [34], setting the parameter “prealigned” to “False” and the parameter “binsize” to “1” to retrieve information necessary for aligning the image with preprocessed spatial coordinates. Subsequently, we applied the “spateo.segmentation.refine alignment” function, specifying “mode”=“rigid”, to align the image accurately. For the human breast cancer dataset from Xenium, we used the affine matrix provided by Xenium to transform and align the image with the spatial coordinates. This was accomplished using the “skimage.transform.AffineTransform” and “skimage.transform.warp” functions from the scikit-image package. Images from the non-small-cell lung cancer dataset (CosMx), the human colon cancer dataset (Visium HD), and the human intestine (CODEX) did not require additional processing.

### Image feature processing

The image patches were fed into a pre-trained ResNet model [35] to extract image features. Specifically, we employed the ResNet18 model, pre-trained on morphological image datasets [36], and removed the last fully connected layer to obtain image feature embeddings. Since stDyer-image was trained using a mini-batch strategy, K-nearest image neighbors were identified for each unit within the same batch based on the cosine distance metric. These image neighbors were then incorporated into the computation of image-related loss during training (see **??**).

### The network structure of stDyer-image and associated probabilistic models

The network architecture of stDyer-image and its associated probabilistic models mirror those of stDyer and are briefly described here. The GMVAE network [13] consists of two main components: an encoder, which models the inference process, and a decoder, which models the generative process. The encoder is a graph attention network [14] that dynamically aggregates gene expression profiles or protein abundance profiles from the neighbors of a given unit in a graph. The decoder reconstructs the gene expression profiles or protein abundance profiles of the target unit. The generative process, parameterized by *θ*, in stDyer-image is represented as follows:

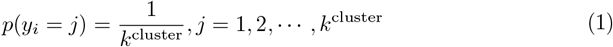

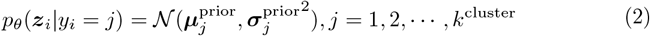

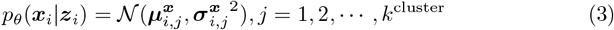

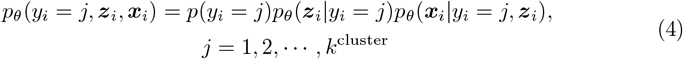

Let ***x***_*i*_, *y*_*i*_ and ***z***_*i*_ denote the gene expression profile or protein abundance profile, cluster label, and latent embedding from the *j*th mode of GMM for the target unit *i*, respectively. The parameters 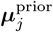 and 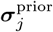 represent the mean and standard deviation of the prior distribution for the *j*th Gaussian mixture. Similarly, 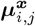 and 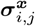 denote the mean and standard deviation of the output distribution (i.e., multivariate Gaussian distribution). Using a multivariate Gaussian distribution as the output distribution stabilizes the training process by avoiding the direct reconstruction of raw gene expression counts or protein abundance profiles. Furthermore, using a continuous distribution enables stDyer-image to adapt to inputs that require normalization. Notably, the output distribution is unit-wise, meaning that each unit has its own unique multivariate Gaussian distribution. This approach is different from other methods that utilize a single negative binomial distribution to represent all gene expression values of each unit. Consequently, the output distribution does not influence the sparsity of the reconstructed values but instead defines the output values as continuous.

The inference process, parameterized by *ϕ*, is formulated as follows:

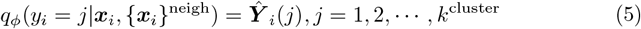

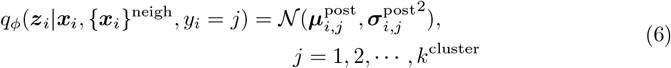

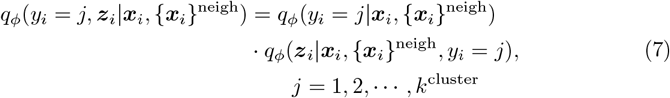

where ***Ŷ*** _*i*_(*j*) denotes the predicted probability that the target unit *i* belongs to the *j*th mode of GMM, and {***x***_*i*_}^neigh^ represents the gene expression profiles or protein abundance profiles of the neighbors of unit *i*. The cluster assignment vector ***Ŷ*** _*i*_(*j*) is inferred using the GAT. 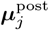 and 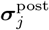 denote the mean and standard deviation of the posterior distribution for the *j*th Gaussian mixture.

### The objective function of stDyer-image

The objective function of stDyer-image consists of an image-related objective function and the original objective function of stDyer. The image-related objective function encourages units with similar image embeddings to share similar soft cluster label. On the other hand, the objective function of stDyer encourages neighboring units to have similar embeddings and cluster labels based solely on gene expression or protein abundance data combined with spatial coordinates. This is achieved by maximizing the log-likelihood of the marginal distribution used to model the gene expression or protein abundance data.

The objective function of stDyer-image can be written as:

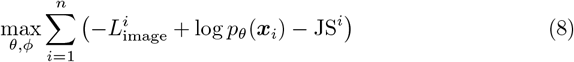

where *L*_image_ is the image-related loss function of unit *i*, ***x***_*i*_ refers to the gene expression profiles or protein abundance profile of unit *i* and JS^*i*^ denotes the Jensen-Shannon divergence of unit *i*.

The image-related loss function is computed for each unit *i* with its *k* image neighbors (default *k*=8) within the mini-batch. These image neighbors are the units whose image embeddings, derived from their associated image patches, are most similar to that of unit *i*. Unit *i* and its image neighbors are encouraged to have higher probabilities for their corresponding soft labels.

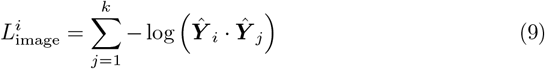

### Integrated gradient analysis

We incorporated the integrated gradient analysis [37] in stDyer-image to identify SVGs. Briefly, it is a technique to estimate the importance of a gene for a model to predict the cluster label of a unit. We followed the same procedure as stDyer and the details of integrated gradient analysis can be referred to [15].

### Evaluation metrics

We used the Adjusted Rand Index (ARI) to evaluate the performance of clustering results [38]. The ARI score measures the agreement between the ground truth cluster labels and the predicted cluster labels, with scores ranging from -1 to 1. For evaluation, all methods are required to predict the same number of clusters as the ground truth. A higher ARI score indicates better clustering performance. Given the ground truth cluster ***A*** = (*a*_1_, *a*_2_, …, *a*_*t*_) and the predicted cluster ***B*** = (*b*_1_, *b*_2_, …, *b*_*p*_), the ARI is defined as follows:

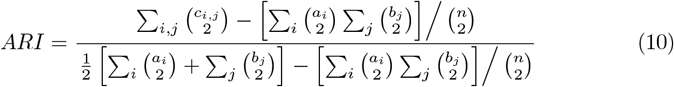

where *c*_*i,j*_ = |*a*_*i*_*∩ b*_*j*_|, and *n* is the number of units.

The Silhouette score [39] is used to obtain the optimal cluster number for the human colon cancer dataset from 10x Visium HD technology. A higher Silhouette score indicates better separateness, while a lower Silhouette score indicates worse separateness. The Silhouette score is computed as the average Silhouette coefficients of all units. The Silhouette coefficient for a unit is computed as follows:

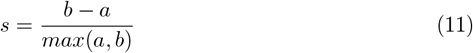

where *a* is the average distance between a unit and all other units in the same cluster, and *b* is the average distance between a unit and all other units in the nearest cluster.

## Availability of data and materials

The NSCLC dataset [1] from CosMx technology is available in https://staging.nanostring.com/products/cosmx-spatial-molecular-imager/ffpe-dataset/nsclc-ffpe-dataset/. The expression profiles and image of mouse brain dataset [2] from Stereo-seq technology can be directly downloaded from https://ftp.cngb.org/pub/SciRAID/stomics/STDS0000058/stomics/Mouse_brain.h5ad and https://ftp.cngb.org/pub/SciRAID/stomics/STDS0000058/Image/Mouse_brain_Adult.tif, respectively. The human breast cancer dataset [3] from 10x Xenium technology was downloaded from https://www.10xgenomics.com/products/xenium-in-situ/preview-dataset-human-breast. The human colon cancer dataset [4] from 10x Visium HD technology was obtained from https://www.10xgenomics.com/cn/datasets/visium-hd-cytassist-gene-expression-libraries-of-human-crc. The SRP data of the human intestine dataset [5] from CODEX technology are accessible at https://datadryad.org/stash/dataset/doi:10.5061/dryad.76hdr7t1p. The corresponding annotation can be accessed at https://datadryad.org/stash/dataset/doi:10.5061/dryad.pk0p2ngrf. The source code of stDyer-image is publicly available at GitHub: https://github.com/ericcombiolab/stDyer-image.

## Authors’ contributions

L.Z. and X.M.Z. conceived and supervised the project. K.X. designed the method, conducted the experiments, and wrote the manuscript. All authors reviewed and approved the manuscript.

## References

[1] He, S., Bhatt, R., Brown, C., Brown, E.A., Buhr, D.L., Chantranuvatana, K., Danaher, P., Dunaway, D., Garrison, R.G., Geiss, G., Gregory, M.T., Hoang, M.L., Khafizov, R., Killingbeck, E.E., Kim, D., Kim, T.K., Kim, Y., Klock, A., Korukonda, M., Kutchma, A., Lewis, Z.R., Liang, Y., Nelson, J.S., Ong, G.T., Perillo, E.P., Phan, J.C., Phan-Everson, T., Piazza, E., Rane, T., Reitz, Z., Rhodes, M., Rosenbloom, A., Ross, D., Sato, H., Wardhani, A.W., Williams-Wietzikoski, C.A., Wu, L., Beechem, J.M.: High-plex imaging of rna and proteins at subcellular resolution in fixed tissue by spatial molecular imaging. Nature Biotechnology 40(12), 1794–1806 (2022) 10.1038/s41587-022-01483-z

[2] Chen, A., Liao, S., Cheng, M., Ma, K., Wu, L., Lai, Y., Qiu, X., Yang, J., Xu, J., Hao, S., Wang, X., Lu, H., Chen, X., Liu, X., Huang, X., Li, Z., Hong, Y., Jiang, Y., Peng, J., Liu, S., Shen, M., Liu, C., Li, Q., Yuan, Y., Wei, X., Zheng, H., Feng, W., Wang, Z., Liu, Y., Wang, Z., Yang, Y., Xiang, H., Han, L., Qin, B., Guo, P., Lai, G., Muoz-Cnoves, P., Maxwell, P.H., Thiery, J.P., Wu, Q.-F., Zhao, F., Chen, B., Li, M., Dai, X., Wang, S., Kuang, H., Hui, J., Wang, L., Fei, J.-F., Wang, O., Wei, X., Lu, H., Wang, B., Liu, S., Gu, Y., Ni, M., Zhang, W., Mu, F., Yin, Y., Yang, H., Lisby, M., Cornall, R.J., Mulder, J., Uhln, M., Esteban, M.A., Li, Y., Liu, L., Xu, X., Wang, J.: Spatiotemporal transcriptomic atlas of mouse organogenesis using dna nanoball-patterned arrays. Cell 0(0) (2022) 10.1016/j.cell.2022.04.003

[3] Janesick, A., Shelansky, R., Gottscho, A.D., Wagner, F., Williams, S.R., Rouault, M., Beliakoff, G., Morrison, C.A., Oliveira, M.F., Sicherman, J.T., Kohlway, A., Abousoud, J., Drennon, T.Y., Mohabbat, S.H., Taylor, S.E.B.: High resolution mapping of the tumor microenvironment using integrated single-cell, spatial and in situ analysis. Nature Communications 14(1), 8353 (2023) 10.1038/s41467-023-43458-x

[4] Oliveira, M.F., Romero, J.P., Chung, M., Williams, S., Gottscho, A.D., Gupta, A., Pilipauskas, S.E., Mohabbat, S., Raman, N., Sukovich, D., Patterson, D., Taylor, S.E.B.: Characterization of immune cell populations in the tumor microenvi-ronment of colorectal cancer using high definition spatial profiling. bioRxiv, 2024–0604597233 (2024) 10.1101/2024.06.04.597233

[5] Hickey, J.W., Becker, W.R., Nevins, S.A., Horning, A., Perez, A.E., Zhu, C., Zhu, B., Wei, B., Chiu, R., Chen, D.C., Cotter, D.L., Esplin, E.D., Weimer, A.K., Caraccio, C., Venkataraaman, V., Schrch, C.M., Black, S., Brbi, M., Cao, K., Chen, S., Zhang, W., Monte, E., Zhang, N.R., Ma, Z., Leskovec, J., Zhang, Z., Lin, S., Longacre, T., Plevritis, S.K., Lin, Y., Nolan, G.P., Greenleaf, W.J., Snyder, M.: Organization of the human intestine at single-cell resolution. Nature 619(7970), 572–584 (2023) 10.1038/s41586-023-05915-x

[6] Zhao, E., Stone, M.R., Ren, X., Guenthoer, J., Smythe, K.S., Pulliam, T., Williams, S.R., Uytingco, C.R., Taylor, S.E.B., Nghiem, P., Bielas, J.H., Gottardo, R.: Spatial transcriptomics at subspot resolution with bayesspace. Nature Biotechnology, 1–10 (2021) 10.1038/s41587-021-00935-2

[7] Varrone, M., Tavernari, D., Santamaria-Martnez, A., Walsh, L.A., Ciriello, G.: Cellcharter reveals spatial cell niches associated with tissue remodeling and cell plasticity. Nature Genetics 56(1), 74–84 (2024) 10.1038/s41588-023-01588-4

[8] Hu, J., Li, X., Coleman, K., Schroeder, A., Ma, N., Irwin, D.J., Lee, E.B., Shinohara, R.T., Li, M.: Spagcn: Integrating gene expression, spatial location and histology to identify spatial domains and spatially variable genes by graph convolutional network. Nature Methods, 1–10 (2021) 10.1038/s41592-021-01255-8

[9] Pham, D., Tan, X., Balderson, B., Xu, J., Grice, L.F., Yoon, S., Willis, E.F., Tran, M., Lam, P.Y., Raghubar, A., Kalita-de Croft, P., Lakhani, S., Vukovic, J., Ruitenberg, M.J., Nguyen, Q.H.: Robust mapping of spatiotemporal trajectories and cell-cell interactions in healthy and diseased tissues. Nature Communications 14(1), 7739 (2023) 10.1038/s41467-023-43120-6

[10] Tang, Z., Li, Z., Hou, T., Zhang, T., Yang, B., Su, J., Song, Q.: Sigra: single-cell spatial elucidation through an image-augmented graph transformer. Nature Communications 14(1), 5618 (2023) 10.1038/s41467-023-41437-w

[11] Xu, C., Jin, X., Wei, S., Wang, P., Luo, M., Xu, Z., Yang, W., Cai, Y., Xiao, L., Lin, X., Liu, H., Cheng, R., Pang, F., Chen, R., Su, X., Hu, Y., Wang, G., Jiang, Q.: Deepst: identifying spatial domains in spatial transcriptomics by deep learning. Nucleic Acids Research 50(22), 131 (2022) 10.1093/nar/gkac901

[12] Bao, F., Deng, Y., Wan, S., Shen, S.Q., Wang, B., Dai, Q., Altschuler, S.J., Wu, L.F.: Integrative spatial analysis of cell morphologies and transcriptional states with muse. Nature Biotechnology 40(8), 1200–1209 (2022) 10.1038/s41587-022-01251-z

[13] Dilokthanakul, N., Mediano, P.A.M., Garnelo, M., Lee, M.C.H., Salimbeni, H., Arulkumaran, K., Shanahan, M.: Deep Unsupervised Clustering with Gaussian Mixture Variational Autoencoders (2016). https://arxiv.org/pdf/1611.02648

[14] Petar Velikovi, Guillem Cucurull, Arantxa Casanova, Adriana Romero, Pietro Li, Yoshua Bengio: Graph attention networks. International Conference on Learning Representations (2022)

[15] Xu, K., Xu, Y., Wang, Z., Zhou, X., Zhang, L.: stdyer enables spatial domain clustering with dynamic graph embedding. bioRxiv, 2024–0508593252 (2024) 10.1101/2024.05.08.593252

[16] Pak, M.G., Shin, D.H., Lee, C.H., Lee, M.K.: Significance of epcam and trop2 expression in non-small cell lung cancer. World Journal of Surgical Oncology 10(1), 53 (2012) 10.1186/1477-7819-10-53

[17] Minghui WANG, Lin FENG, Ping LI, Naijun HAN, Yanning GAO, Ting XIAO: Hsp90ab1 protein is overexpressed in non-small cell lung cancer tissues and associated with poor prognosis in lung adenocarcinoma patients. Chinese Journal of Lung Cancer 19(2) (2016)

[18] Cords, L., Tietscher, S., Anzeneder, T., Langwieder, C., Rees, M., Souza, N., Bodenmiller, B.: Cancer-associated fibroblast classification in single-cell and spatial proteomics data. Nature Communications 14(1), 4294 (2023) 10.1038/s41467-023-39762-1

[19] Iwano, T., Masuda, A., Kiyonari, H., Enomoto, H., Matsuzaki, F.: Prox1 post-mitotically defines dentate gyrus cells by specifying granule cell identity over ca3 pyramidal cell fate in the hippocampus. Development 139(16), 3051–3062 (2012) 10.1242/dev.080002

[20] Lavado, A., Lagutin, O.V., Chow, L.M.L., Baker, S.J., Oliver, G.: Prox1 is required for granule cell maturation and intermediate progenitor maintenance during brain neurogenesis. PLOS Biology 8(8), 1000460 (2010) 10.1371/journal.pbio.1000460

[21] Iijima, T., Miura, E., Watanabe, M., Yuzaki, M.: Distinct expression of c1q-like family mrnas in mouse brain and biochemical characterization of their encoded proteins. European Journal of Neuroscience 31(9), 1606–1615 (2010) 10.1111/j.1460-9568.2010.07202.x

[22] Fitzgerald, N.D., Day, J.J.: Neuronal heterogeneity in the ventral tegmental area: Distinct contributions to reward circuitry and motivated behavior. Addiction Neuroscience 14, 100191 (2025) 10.1016/j.addicn.2024.100191

[23] Vanauberg, D., Schulz, C., Lefebvre, T.: Involvement of the pro-oncogenic enzyme fatty acid synthase in the hallmarks of cancer: a promising target in anti-cancer therapies. Oncogenesis 12(1), 16 (2023) 10.1038/s41389-023-00460-8

[24] Xu, S., Chen, T., Dong, L., Li, T., Xue, H., Gao, B., Ding, X., Wang, H., Li, H.: Fatty acid synthase promotes breast cancer metastasis by mediating changes in fatty acid metabolism. Oncology Letters 21(1), 27 (2021) 10.3892/ol.2020.12288

[25] Sun, J., Huang, J., Lan, J., Zhou, K., Gao, Y., Song, Z., Deng, Y., Liu, L., Dong, Y., Liu, X.: Overexpression of cenpf correlates with poor prognosis and tumor bone metastasis in breast cancer. Cancer Cell International 19(1), 264 (2019) 10.1186/s12935-019-0986-8

[26] O’Brien, S.L., Fagan, A., Fox, E.J.P., Millikan, R.C., Culhane, A.C., Brennan, D.J., McCann, A.H., Hegarty, S., Moyna, S., Duffy, M.J., Higgins, D.G., Jirstrm, K., Landberg, G., Gallagher, W.M.: Cenp-f expression is associated with poor prognosis and chromosomal instability in patients with primary breast cancer. International Journal of Cancer 120(7), 1434–1443 (2007) 10.1002/ijc.22413

[27] Ehrencrona, E., van der Post, S., Gallego, P., Recktenwald, C.V., Rodriguez-Pineiro, A.M., Garcia-Bonete, M.-J., Trillo-Muyo, S., Bckstrm, M., Hansson, G.C., Johansson, M.E.V.: The iggfc-binding protein fcgbp is secreted with all gdph sequences cleaved but maintained by interfragment disulfide bonds. The Journal of biological chemistry 297(1), 100871 (2021) 10.1016/j.jbc.2021.100871

[28] Castro, C.D., Flajnik, M.F.: Putting j chain back on the map: how might its expression define plasma cell development? The Journal of Immunology 193(7), 3248–3255 (2014) 10.4049/jimmunol.1400531

[29] Ma, B., Li, F., Ma, B.: Down-regulation of col1a1 inhibits tumor-associated fibroblast activation and mediates matrix remodeling in the tumor microenvi-ronment of breast cancer. Open Life Sciences 18(1), 20220776 (2023) 10.1515/biol-2022-0776

[30] Jia, S.-N., Han, Y.-B., Yang, R., Yang, Z.-C.: Chemokines in colon cancer progression. Seminars in Cancer Biology 86(Pt 3), 400–407 (2022) 10.1016/j.semcancer.2022.02.007

[31] Zhao, Q., Zong, H., Zhu, P., Su, C., Tang, W., Chen, Z., Jin, S.: Crosstalk between colorectal cscs and immune cells in tumorigenesis, and strategies for targeting colorectal cscs. Experimental Hematology & Oncology 13(1), 6 (2024) 10.1186/s40164-024-00474-x

[32] Wolf, F.A., Angerer, P., Theis, F.J.: Scanpy: large-scale single-cell gene expres-sion data analysis. Genome Biology 19(1), 15 (2018) 10.1186/s13059-017-1382-0

[33] Charles R. Harris, K. Jarrod Millman, Stfan J. van der Walt, Ralf Gommers, Pauli Virtanen, David Cournapeau, Eric Wieser, Julian Taylor, Sebastian Berg, Nathaniel J. Smith, Robert Kern, Matti Picus, Stephan Hoyer, Marten H. van Kerkwijk, Matthew Brett, Allan Haldane, Jaime Fernndez del Ro, Mark Wiebe, Pearu Peterson, Pierre Grard-Marchant, Kevin Sheppard, Tyler Reddy, Warren Weckesser, Hameer Abbasi, Christoph Gohlke, Travis E. Oliphant: Array programming with numpy. Nature 585(7825), 357–362 (2020) 10.1038/s41586-020-2649-2

[34] Qiu, X., Zhu, D.Y., Lu, Y., Yao, J., Jing, Z., Min, K.H., Cheng, M., Pan, H., Zuo, L., King, S., Fang, Q., Zheng, H., Wang, M., Wang, S., Zhang, Q., Yu, S., Liao, S., Liu, C., Wu, X., Lai, Y., Hao, S., Zhang, Z., Wu, L., Zhang, Y., Li, M., Tu, Z., Lin, J., Yang, Z., Li, Y., Gu, Y., Ellison, D., Chen, A., Liu, L., Weissman, J.S., Ma, J., Xu, X., Liu, S., Bai, Y.: Spatiotemporal modeling of molecular holograms. Cell 0(0) (2024) 10.1016/j.cell.2024.10.011

[35] He, K., Zhang, X., Ren, S., Sun, J.: Deep residual learning for image recognition. In: Proceedings of the IEEE Conference on Computer Vision and Pattern Recognition (CVPR), pp. 770–778 (2016). https://openaccess.thecvf.com/content_cvpr_2016/html/HeDeepResidualLearningCVPR2016paper.html

[36] Ciga, O., Xu, T., Martel, A.L.: Self supervised contrastive learning for digital histopathology. Machine Learning with Applications 7, 100198 (2022) 10.1016/j.mlwa.2021.100198

[37] Sundararajan, M., Taly, A., Yan, Q.: Axiomatic attribution for deep networks. In: Precup, D., Teh, Y.W. (eds.) Proceedings of the 34th International Conference on Machine Learning. Proceedings of Machine Learning Research, vol. 70, pp. 3319–3328. PMLR, New York, New York, USA (2017). https://proceedings.mlr.press/v70/sundararajan17a.html

[38] Hubert, L., Arabie, P.: Comparing partitions. Journal of Classification 2(1), 193– 218 (1985) 10.1007/BF01908075

[39] Rousseeuw, P.J.: Silhouettes: A graphical aid to the interpretation and validation of cluster analysis. Journal of Computational and Applied Mathematics 20, 53–65 (1987) 10.1016/0377-0427(87)90125-7

